# Extended longevity geometrically-inverted proximal tubule organoids for protein uptake studies

**DOI:** 10.1101/2022.03.24.485493

**Authors:** Eric Parigoris, Ji-Hoon Lee, Amy Yunfan Liu, Xueying Zhao, Shuichi Takayama

**Affiliations:** Wallace H. Coulter Department of Biomedical Engineering, Georgia Institute of Technology, Atlanta, GA, United States; The Parker H. Petit Institute of Bioengineering and Bioscience, Georgia Institute of Technology, Atlanta, GA, United States; Department of Physiology, Morehouse School of Medicine, Atlanta, GA, United States

**Keywords:** Organoids, kidney, proteinuria, basal-in/apical-out

## Abstract

While some *in vitro* platforms have been adapted to study proteinuric kidney disease, organoids have been challenging to study the disease. This is because apical access is historically difficult, and this is the surface on which megalin (LRP2), an endocytic receptor responsible for tubular reabsorption of filtered plasma proteins, resides. Based on our previous geometrically-inverted organoids, this study established high-throughput basal-in and apical-out proximal tubule organoids to study proteinuric kidney disease in a more physiologically consistent manner. Organoids successfully formed around a minimal Matrigel scaffold, and were maintained in culture for 90+ days, the longest reported hanging drop culture to date. The proximal tubule organoids exhibited good polarization, showed upregulation of maturity markers, such as aquaporin-1 and megalin, and experienced less epithelial-to-mesenchymal (EMT) transition compared to 2D cells. To assess protein uptake, fluorescent albumin was placed in the surrounding media, facing the apical surface, and organoids demonstrated functional protein uptake even at 90 days. To mimic proteinuric conditions, organoids were exposed to human serum albumin and released kidney injury molecule-1 (KIM-1), a common biomarker for kidney injury, in both dose- and time-dependent manners. While this study focuses on applications for modeling proteinuric kidney disease conditions, these organoids are envisioned to have broad utility where apical proximal tubule cell access is required.

## 1. Introduction

Approximately 37 million people in the United States, or 15% of the population, are living with chronic kidney disease (CKD) [1], with a large portion of these Americans being undiagnosed. One of the main determinants of CKD is proteinuria, or high levels of protein in the urine; this is typically caused by juxtaglomerular imbalance in which too much protein can pass through the selective glomerular barrier. In addition to CKD, proteinuria can be caused by a variety of conditions, ranging from temporary ones, such as dehydration and intense exercise, to underlying conditions, such as diabetes, hypertension, or nephrotic syndrome [2, 3]. While imbalances in the glomerulus, or the selective filtration barrier of the nephron (Figure 1a), are the physiological cause of proteinuria, the consequences are seen in the proximal tubule. This is because a majority of protein reabsorption occurs in the proximal tubule, causing it to be the major site of tubular injury. The loop of Henle, distal tubule, and collecting duct do not significantly contribute to protein reabsorption, but are rather focused on water and ion reabsorption (Figure 1a).

**Figure 1.**
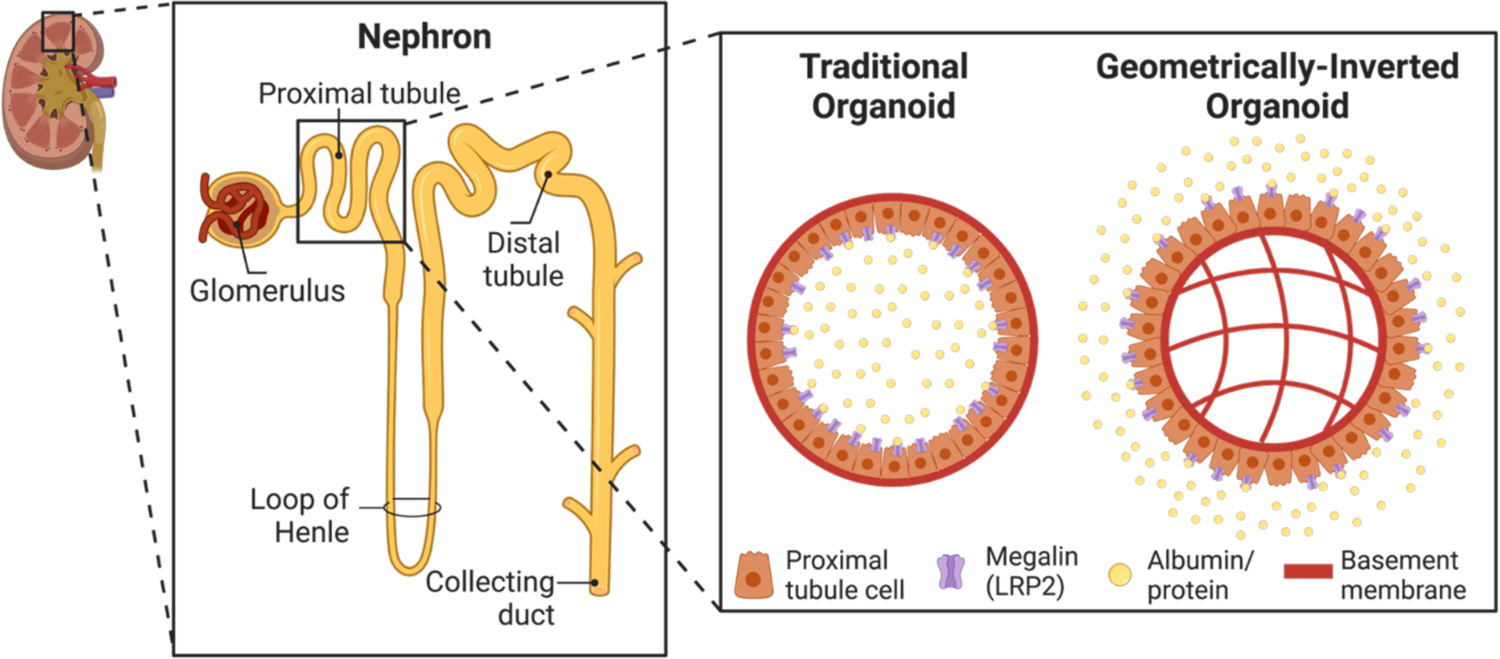
Establishment of geometrically-inverted organoids. Schematic of a nephron, the functional unit of the kidney. Inset shows a comparison of typical organoids, and geometrically-inverted organoids. Since megalin is located on the apical surface of proximal tubule cells, apical-out organoids allow for more ease of access of megalin and therefore proteinuria studies.

Previous studies have shown that high albumin levels in the urinary space cause elevation in damage signals, which are not present in significant amounts in the absence of injury. A common one of particular interest is kidney injury molecule-1 (KIM-1), an early biomarker of proximal tubule injury, due to its easy and non-invasive detection [4]. Increased KIM-1 expression may enhance albumin uptake via receptor-mediated endocytosis on the apical side of the proximal tubule cells [5]. This in turn, can exacerbate the deleterious effects of proteinuria, leading to cell injury, inflammation, and cell apoptosis [3].

While animal experiments are effective for proteinuria studies, there are known organism-specific differences in important transporters, and they pose ethical and cost concerns [6, 7]. To study such a disease, many have turned to *in vitro* platforms such as 2D cell cultures, Transwells, organ-on-a-chip systems, and organoids to minimize the cost and challenges associated with animal models; they are gradually becoming adopted, as they have improved our understanding of normal kidney function, diseased kidney states, and drug-induced nephrotoxicity [8, 9]. While 2D cultures are high-throughput and well standardized, they lack physiological relevance and cannot recapitulate the 3D architecture found in the body. They also cannot survive long-term culture, and growth on stiff substrates induce epithelial-to-mesenchymal transition (EMT) [10]. On the other hand, kidney-on-a-chip systems facilitate exposure of fluid shear stress (FSS) to the apical surface of proximal tubule cells and therefore promote enhanced polarization, barrier function, and overall cell maturity [4]. Kidney-on-a-chip systems also generally lack the 3D architecture found *in vivo* and their implementation can be low-throughput, technically demanding, and lack standardization [4,8,9] (Supplementary Figure 1).

Most 3D kidney organoid models are derived from induced pluripotent stem cells (iPSCs), embryonic stem cells (ESCs) [11–14], or adult stem cells [15, 16], and only a few 3D proximal tubule-specific models have been successfully developed [10,17,18]. Many 3D models can be grown in long-term culture, and models have shown that they have upregulated megalin, an important indicator of cellular differentiation and maturation [17, 18]. This is also a major advantage for proteinuria studies, as this is the receptor responsible for albumin interactions and reabsorption. Traditional organoids, however, have inaccessible lumens or apical surfaces, which is a main consideration for the application of the pathophysiological of proteinuria, in which high protein levels are exposed to the apical side of the proximal tubule [8]. Access to the apical surface is therefore critical for studying proteinuria in 3D organotypic systems.

There has been recent attention given to geometrically-inverted organoids, in which the basolateral and apical sides are reversed [19–22]. Our lab has established basal-in cultures for breast organoids grown from MCF10A cells [19, 20]. These organoids, grown in a one-drop, one-organoid hanging drop culture or ultra-low attachment plates (ULA) have proven to be more reproducible and uniform in size compared to other organoid models [20, 23]. Other geometrically-inverted organotypic structures have been reported for the intestine [21, 24], lung [22], and kidney [25–29]. Of interest, many have studied kidney cysts made of distal tubule cells (MDCK) that have reversed polarity. To our knowledge, there have been no attempts to generate proximal tubule organoids with reversed polarity, in which the basolateral side faces inward, and the apical side is outward-facing. Because of the ease of apical access, this platform would facilitate its use in proteinuria applications, thereby bypassing the physiological inconsistencies with 2D culture, and avoiding cost and ethical concerns associated with animal studies This study therefore developed a high-throughput proximal tubule organoid platform that is geometrically-inverted, in which the basal side is inwards facing and the apical side is outwards facing (Figure 1). Material considerations were given to optimize organoid formation around a Matrigel core, and the organoids were subsequently maintained in hanging drop culture for 90+ days. The organoids exhibited basal-in and apical-out polarity, as indicated by Na^+^K^+^ATPase and laminin-5 localized to the organoid interior (basolateral side) and phalloidin and ezrin/villin-2 localized to the organoid exterior (apical side). Immunostaining and bulk RNA sequencing (RNAseq) revealed that the organoids matured over time, exhibited less EMT compared to 2D samples, and consistently expressed megalin, an important receptor for protein uptake. We therefore utilized these organoids to mimic proximal tubule injury due to high protein concentrations exposed to the apical surface. After confirming apical protein uptake of fluorescein isothiocyanate bovine serum albumin (FITC-BSA), we showed both dose dependent and time dependent responses of KIM-1 production to incubation with human serum albumin (HSA).

## 2. Materials and methods

### 2.1. 2D culture

Human renal proximal tubule epithelial cells (RPTECs) were grown in renal epithelial cell basal media (ATCC) supplemented with 0.5% fetal bovine serum (FBS), 10 nM triiodothyronine, 10 ng/mL epidermal growth factor (EGF), 100 ng/mL hydrocortisone hemisuccinate, 5 ug/mL insulin, 1.0 µM epinipherine, 5 µg/mL transferrin, 2.4 mM L-glutamine, and 1% penicillin-streptomycin (pen-strep). RPTECs were maintained in T75 flasks and passaged at 95% confluency. Primary RPTECs were used until passage 6. All experiments were performed with lot 70001234, in which the donor was a 56-year-old Caucasian male.

### 2.2. 3D organoid culture

RPTEC organoids were maintained in hanging drop culture as previously described [20, 23]. Briefly, custom hanging drop plates were soaked overnight in 0.1% Pluronic solution (Sigma, #542342). Plates were rinsed with water and sterilized using a UVP cross linker (Analytik Jena). Hanging drop plates were sandwiched between a 96-well round bottom plate. To prevent droplet evaporation, gauze pad and sterile water with 1% pen-strep were added to the side troughs, and 150 μL deionized water + 1% pen-strep was added to each well of the 96-well plate.

The RPTEC seeding solution consisted of 120,000 RPTEC cells/mL, 120 μg/mL Matrigel, 0.24% methylcellulose (Methocel), and 2.4% FBS. To ensure a basal-in and apical-out phenotype and proper encapsulation of Matrigel in the organoid core, cold Matrigel was added to prewarmed media, Methocel (A4M, Sigma), and FBS (Gemini). Upon the addition of Matrigel (Corning, #356231), the solution was mixed thoroughly, cells were immediately added to the solution, and 25 μL (corresponding to 3000 cells/droplet) was added to each droplet of the plate using a multi-channel repeater pipette. Hanging drop plates were incubated at 37°C and 5% CO_2_. After 72 hours, 3 consecutive media changes were performed using the Cybio FeliX liquid handler (Analytik Jena), each removing 9 μL and adding 10 μL. Media changes were then performed every 2-3 days, with 2 consecutive rounds of 9 μL removal and 9 μL addition.

Because the plates were maintained for 90+ days in culture, evaporation of the droplets needed to be carefully considered. To account for this, 5 μL of additional media was added, particularly to the outer droplets. The deionized water was replenished in the bottom receiver plate as needed. During routine media changes, the amount added or removed was adjusted accordingly based on the size of the droplets; if they looked too big, a little media was removed and if they looked too small, additional media was added.

### 2.3. Live-organoid imaging and morphology quantification

RPTEC organoids were imaged every 2-3 days with a EVOS FL 2 Auto microscope (Thermo Fisher) with a 4x objective. To quantify the area, diameter, and roundness of the organoids, raw image files were imported into ImageJ [30]. The images were converted to binary, inverted, and holes were filled. The “Analyze Particles” feature was used to calculate the area and roundness. To determine the mean diameter, the organoid was assumed as a perfect circle and the diameter was calculated.

### 2.4. Histology, immunofluorescence staining, and microscopy

RPTEC organoid histology, staining, and imaging were performed based on our previously described protocols [19, 20]. Organoids were harvested from the hanging drop plate, washed with phosphate-buffered saline (PBS), and fixed for 30 minutes at room temperature in 4% paraformaldehyde (Alfa Aesar). Organoids were washed again with PBS, and stained for 10 minutes in 0.5% methylene blue (Ricca Chemical Company) in PBS to aid in visualization during the sectioning process. Excess dye was then washed out, and the samples were transferred to a block and filled with optimal cutting temperature (OCT, Tissue-Tek). Isopentane (Sigma) was then cooled in liquid nitrogen, and samples were flash frozen and stored in −80°C until use.

A CryoStar NX70 cryostat (Thermo Fisher) was used to obtain 20 µm thick sections. For hematoxylin and eosin (H&E) staining, organoids were brought to room temperature and stained with an Autostainer XL (Leica). Slides were coverslipped with Xylene and Cytoseal 60 (Richard-Allan Scientific), and imaged in color with a DMi1 microscope (Leica) and 10x air objective. For immunostaining, slides were brought to room temperature and a hydrophobic pen was used to highlight the region containing the organoid section. Samples were washed with PBS, permeabilized with 0.2% triton X-100 (Sigma) in PBS 100 for 5 minutes, and washed again with PBS. Samples were incubated for 1 hour at room temperature with 4% BSA (Millipore Sigma, #82-067) in PBS. The following primary antibodies were prepared in 1% BSA in PBS solutions at the following dilutions: rabbit anti-megalin (Abcam #ab76969, 1:500 dilution), mouse anti-aquaporin-1 (Santa Cruz #sc-32737, 1:200 dilution), mouse anti-ezrin/villin-2 (BD #610602, 1:200 dilution), rabbit anti-Na^+^K^+^ATPase (Abcam #ab76020, 1:250 dilution), and rabbit anti-laminin-5 (Abcam #ab14509, 1:200 dilution). Primary antibodies were added to the samples and incubated at 4°C overnight. Upon removal of the primary antibodies, samples were washed with 1% BSA in PBS, and the following secondary antibody solutions were prepared in 1% BSA: Goat anti-mouse Alexa Fluor 488 (Invitrogen #A-11001, 1:2000 dilution) and goat anti-rabbit Alexa Fluor 594 (Invitrogen #A-11012, 1:1000 dilution). Secondary antibodies were incubated at room temperature for 2 hours along with Alexa Fluor 488 phalloidin (Invitrogen #A12379, 1:40 dilution). Secondary antibodies were removed, samples were washed with PBS, and incubated with DAPI (Thermo Fisher, 1.4 x 10^-6^ uM) for 15 minutes at room temperature. Samples were given a final rinse in PBS and sealed with ProLong Diamond Antifade (Thermo Fisher, #P36965) mounting media. The organoids were imaged using a DMi8 epifluorescence microscope (Leica) equipped with 10x, 20x, and 40x air objectives.

### 2.5. Quantification of organoid cell coverage and percentage of aquaporin-1 (AQP1) and megalin (LRP2) positive cells

To quantify the extent of organoid cell coverage (cells/perimeter), the images from H&E histology sections were utilized. The number of nuclei (only at the organoid periphery), represented by the dark purple spots, were manually counted in ImageJ. The projected cell area was quantified using the methods discussed in section 2.3. The number of cells was divided by the perimeter for each image. Five images were analyzed for days 8, 16, 30, and 90. For the AQP1 and LRP2 quantification, immunofluorescent images were used. For each image, the total number of nuclei were manually counted, and then the number of cells expressing AQP1 and LRP2 were counted. For each marker, the number of positive cells was divided by the total number of cells in the image to calculate the percentage of positive cells. Five images were analyzed for day 16, 30, and 90 organoids.

### 2.6. Bulk RNA sequencing

5-10 organoids per replicate were pooled from hanging drop plates to RNase-free tubes. Samples were prepared for 2D RPTECs (∼10^6^ cells) and day 31 organoids. Organoid media was then removed and 300 μl of RLT lysis buffer with 1% beta-mercaptoethanol was added to each tube. The lysate was vortexed for one minute, and then immediately flash frozen with liquid nitrogen. Frozen lysates were kept in −80°C until further processing. The samples were shipped to the Emory Integrated Genomics Core (EIGC) for RNA extraction, sequencing library preparation, and sequencing. Illumina sequencers and kits were chosen based on the number of samples, availability, and read depths (>20M per sample).

### 2.7. Differential gene expression analysis

FASTQ files provided by EIGC were downloaded to Georgia Tech’s high-performance computing cluster called Partnership for an Advanced Computing Environment (PACE). Quality control was first performed using FastQC. Upon passing quality requirements, the reads were aligned using STAR, or mapped using Salmon. A custom Python script was written to generate a gene count matrix. For Salmon transcripts, a gene count matrix was generated using R. Genes with less than 0.5 CPM (counts per million) in at least 3 samples were filtered out. Differential gene expression analysis was conducted using the EdgeR package. PCA analysis was performed to examine the samples for possible outliers prior to further analysis. The counts were normalized based on library size using the default Trimmed mean of M-values (TMM) method from EdgeR. A comparison was run between the two groups through a generalized linear model (GLM), and likelihood-ratio test (LRT) to identified statistically significant differential expressions. The cutoff for significance was set at under an alpha value of 0.05 for the adjusted p-value, and the log_2_ fold change (LFC) of ±1. A volcano plot comparing 2D and day 31 organoid groups was then generated. The same alpha and LFC cutoffs were used to label up-, down-regulated, and statistically insignificant genes.

### 2.8. Heatmap generation

A custom R script was written to plot heatmaps containing genes-of-interest. Briefly, the script takes in a spreadsheet with gene symbols and their categories. The R package pheatmap was used to compute a z-score matrix, which was then presented in a heatmap.

### 2.9. Protein uptake studies

For albumin uptake studies, we utilized all components of the RPTEC growth media, except 0.5% FBS. FITC-BSA was added to the serum-free growth media at a concentration of 50 μg/mL. 200 μL of media was added to each well of 96-well ultra-low attachment (ULA) plate (Corning, #7007). Organoids were then transferred from the hanging drop plate to an individual well of the ULA plate. Care was taken to introduce as little media from the hanging drop plate to the ULA plate as possible. The ULA plate was incubated at 37°C and 5% CO_2_ for 2 hours. After the incubation, the extracellular FITC-BSA was washed out with PBS or serum-free growth media. To prevent aspiration of the organoids from the bottom of the wells, we performed 10 consecutive half media changes (remove 100 μL, add 100 μL) to remove the excess FITC-BSA. Organoids were then imaged using an in-incubator microscope (Incucyte S3, Sartorius) using brightfield, phase contrast, and a GFP filter.

### 2.10. HSA toxicity and KIM-1 studies

As with the albumin uptake studies, serum-free growth media was utilized, as the serum contains some albumin that could interfere with the experiment. HSA (Sigma, #A4327) was added to the serum-free growth media at the desired concentration, and the solution was sterile filtered. The stock solution was serially diluted using a 1:2 dilution ratio; concentrations ranged from 0.078 mg/mL to 2.5 mg/mL. 200 μL of each solution was transferred to wells in a 96-well ULA plate. Organoids were then transferred from the hanging drop plate to the HSA solutions in the ULA plates. The organoids were incubated for the desired timepoint at 37°C and 5% CO_2_. Organoids were incubated for 48 hours for dose-dependent experiments, and 3, 6, 12, 24, 48, and 72 hours for the time series experiments. At the appropriate timepoint, the supernatant from the wells was collected, flash frozen in liquid nitrogen, and stored at −80°C until needed. To quantify the level of KIM-1 production, and an ELISA kit (Enzo, #ADI-900-226-0001) was utilized. The kit was used according to the manufacturer’s instructions, and a BioTek Synergy H4 microplate reader was used for absorbance readings at 450 nm.

### 2.11. Statistical analysis

To quantify cell area, diameter, and roundness, the same 16 organoids were tracked for 67 days. No statistical analysis was performed. For the quantification of cell coverage, 5 samples were used per condition, and a one-way ANOVA with Tukey’s multiple comparisons was used to analyze the data. To identify differentially expressed genes from the RNAseq data, LRT was performed on each pre-filtered gene. Unadjusted p-values computed from LRT were corrected with the Benjamini-Hochberg procedure. For quantification of percentages of AQP1 and LRP2 positive cells, 5 samples were used for each timepoint. A one-way ANOVA with Tukey’s multiple comparisons was used to analyze the data. For proteinuria experiments, analysis for days 35 and 90 organoids had n=4 samples per condition. A one-way ANOVA with Tukey’s multiple comparisons was applied to the data. For the KIM-1 time course data, n=4 samples were used per condition, and a two-way ANOVA with Šīdák’s multiple comparison tests was utilized. In all graphs, error bars represent the standard deviation. For all statistical tests, * indicates p ≤ 0.05, ** indicates p ≤ 0.01, *** indicates p ≤ 0.001, and **** indicates p ≤ 0.0001.

## 3. Results and Discussion

### 3.1. Minimal Matrigel scaffolding allows for proximal tubule organoid formation

Based on our recent publication of basal-in MCF10A organoids grown in hanging drop culture [20], we tested the ability to extend the methods to human renal proximal tubule epithelial cells (RPTECs). Of interest, these cells are primary kidney proximal tubule cells, compared to previously utilized immortalized cell lines. To optimize high-throughput organoid formation, we tested several different ranges of cell numbers, Matrigel concentrations, temperature conditions, and serum concentrations, many of which led to poor organoid formation (Supplementary Figure 2). We found that a comparable cell number (3000 cells/drop) and Matrigel concentration (110 – 130 μg/mL) to the breast organoids successfully translated to proximal tubule cells to form geometrically-inverted organoids (Figure 2a and 2b). However, the addition of approximately 2% fetal bovine serum (FBS) to the seeding solution yielded the best organoid formation, compared to the 10% FBS added to MCF10A organoids during seeding. It is important to note that RPTECs, even in 2D, are grown in low-serum conditions (0.5% FBS). In line with previous optimizations [20,23,31,32], 0.24% methycellulose (Methocel) was added, and its exclusion resulted in many smaller acinar structures (Supplementary Figure 2). Finally, as with our previous mammary basal-in organoids, the solution temperatures and mixing procedure was critical for proper organoid formation. Methocel, FBS, and growth media were well mixed and thoroughly warmed to 37°C. Immediately before seeding, cold Matrigel was added to the mixture, the cells were added, and the solution was thoroughly mixed before pipetting into hanging drop plates. If the order was varied or if Matrigel was added to room temperature media, we saw improper organoid formation (Supplementary Figure 2).

**Figure 2.**
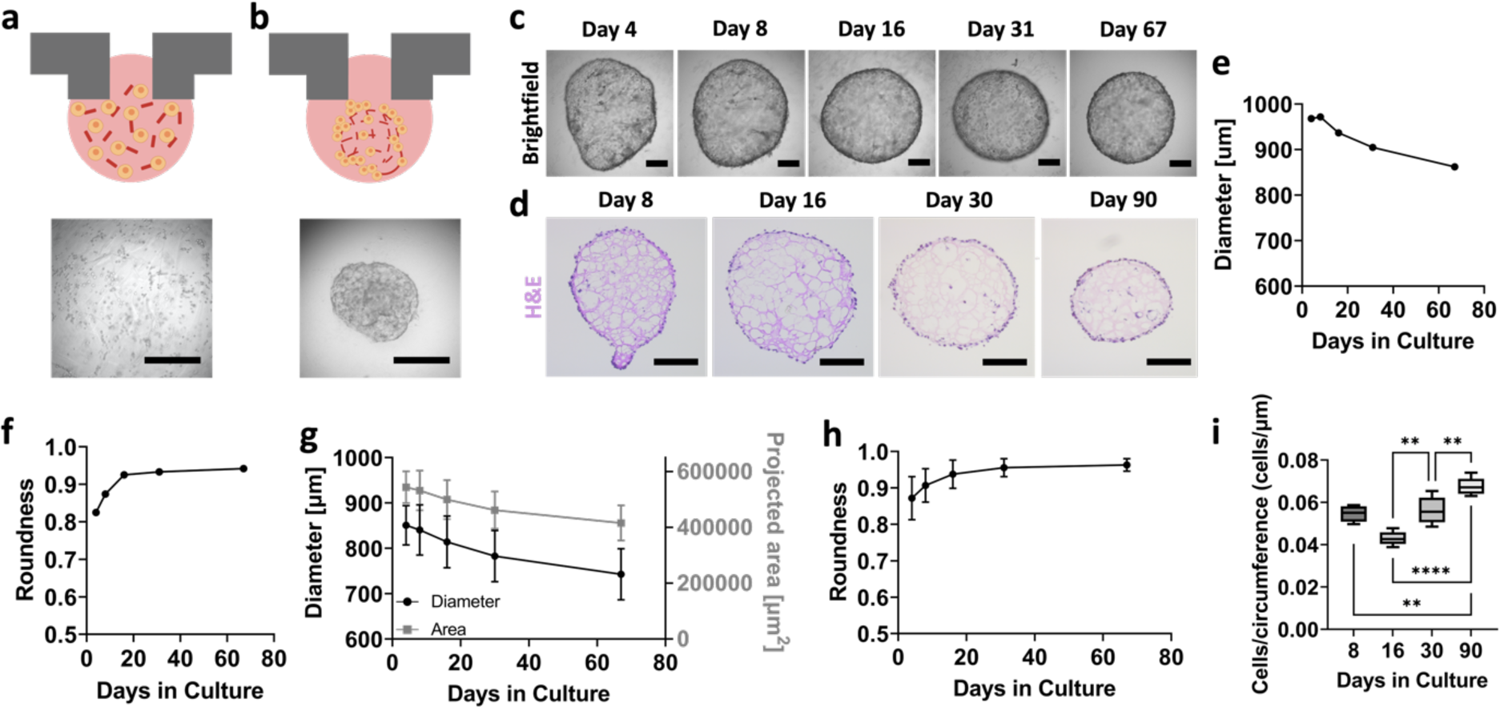
Optimization of geometrically-inverted proximal tubule organoid formation. (a) Schematic and brightfield image of RPTECs at time of seeding, where cells are homogeneously mixed among partially gelled Matrigel. (b) Schematic and brightfield image of organoid formation within 24-48 hours. Scalebars represent 500 µm. (c) Time course brightfield of whole organoids, with the same organoid tracked over time. Scalebars represent 200 µm. (d) H&E images of representative organoid sections at days 8, 16, 30, and 90 of their growth. Scalebars represent 200 µm. (e) Diameter and (f) roundness values for the particular organoid shown in panel (c). (g) Area, diameter, and (h) roundness values for organoids for days 4-67 of their growth. (i) Quantification of cell coverage per circumference length of RPTEC organoids at days 8, 16, 30, and 90 of their growth. For panels (g) and (h), n = 16 per timepoint. For panel (i), n=5 per timepoint. One-way ANOVA was performed using Tukey’s multiple comparisons. ** denotes p ≤ 0.01 and **** denotes p ≤ 0.0001.

We further examined the importance of the minimal Matrigel scaffolding concept. First, we explored the acellular seeding media. We observed that a prewarmed solution composed of Methocel, FBS, and growth media did not have any gelling when pipetted into a hanging drop plate. However, when cold Matrigel was added to the prewarmed media, we observed significant gelling immediately after pipetting into the hanging drop plate (Supplementary Figure 2). Upon the addition of cells, we saw that the partially gelled Matrigel spread throughout the well, and provided a minimal scaffold for the cells to attach to. After 24-48 hours, as with our previous observations, the cells migrated to surround the Matrigel, creating an organoid with a basement-membrane core (Figure 2a and 2b).

Upon successful formation of RPTEC organoids and characterization of the minimal Matrigel scaffold, we maintained them in hanging drop culture for 90+ days, the longest hanging drop culture of any kind, including our previously reported mammary organoids maintained for a maximum of 25 days in hanging drop culture [20]. Live/dead staining revealed that the organoids remained viable at days 65 and 90 of their culture (Supplementary Figure 3). We imaged and tracked organoids every 2-3 days, and representative images of organoids at day 8, 16, 30, and 90 are shown in Figure 2c. Histology and H&E staining was also performed at these timepoints (Figure 2d). Figures 2e and 2f show the morphological features of the particular organoid shown in Figure 2c. Figure 2g shows data from n=16 organoids, and demonstrates a decrease in area/diameter over time, an observation that can also be seen qualitatively in Figures 2c and 2d. Figure 2h shows that their roundness increases over time. These findings were consistent for multiple independent experiments (Supplementary Figure 4).

To quantify cell coverage of the organoid, we manually counted the number of cell nuclei from H&E images, and divided it by the organoid circumference. It is interesting to note that the number of cells per circumference length on the organoids increased between days 16, 30, and 90 of their culture. Although cell numbers were comparable between the conditions, the area decreased over time, especially by day 90 (Figure 2i). Therefore, the organoids experienced an overall increase in cell coverage that is even apparent from the H&E images (Figure 2d). Additional description of extended cultures beyond 90 days is presented in Supplementary Figure 4.

Overall, from a materials perspective, it is necessary to consider the importance of Matrigel in organoid formation and maintenance. From Figure 2d, the network of pink hues suggests that Matrigel is entrapped inside the organoid core. Interestingly, this pink hue fades over time, suggesting the degradation of Matrigel. Although the presence and maintenance of Matrigel in the core is the first indication of basal-in phenotype, we aimed to further explore how this would affect mechanisms for inversion and polarization of RPTEC organoids.

### 3.2. RPTEC organoids exhibit a basal-in and apical-out phenotype

To examine the extent of polarization in the RPTEC organoids, we utilized immunofluorescence staining of relevant apical and basolateral markers. Staining of laminin-5 (red), a basement membrane marker, was localized to the interior side of the organoids at all timepoints. Counterstaining with DAPI revealed that the red is more localized to the interior of the organoid for days 16, 30, and 90 of culture (Figure 3a). Throughout the duration of the organoid culture, laminin-5 exhibited development, maturation, and thickening over time. The thickest region was for day 90, and co-staining with phalloidin (apically located in proximal tubule cells) showed clear localization of the laminin-5 on the organoid interior and phalloidin to the organoid exterior (Figure 3a, inset). A similar trend can be observed when examining co-staining of Na^+^K^+^ATPase, a basolateral marker, and ezrin/villin-2, an apical brush border marker, for days 16, 30, and 90 (Figure 3b). Na^+^K^+^ATPase showed localization to the organoid interior (basolateral side), and ezrin/villin-2 was localized to the organoid exterior (apical side).

**Figure 3.**
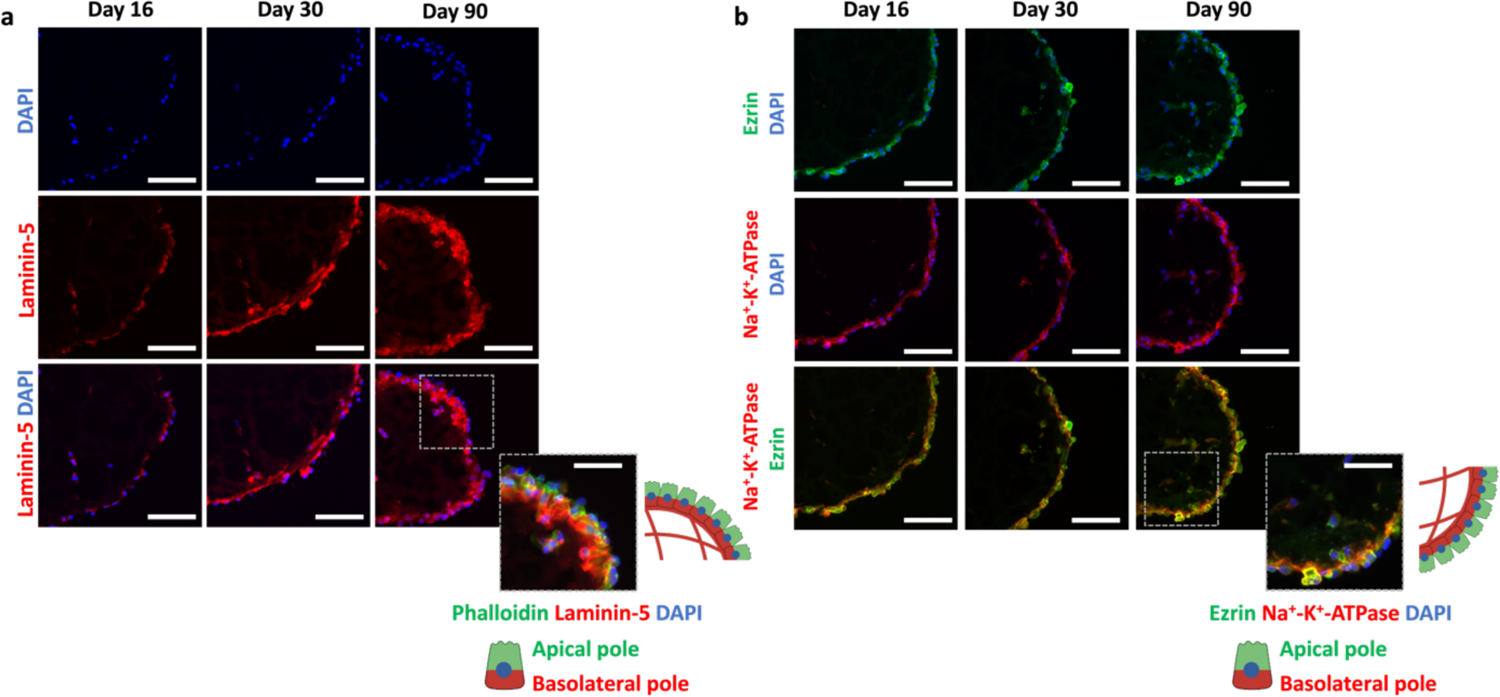
RPTEC organoids exhibit basal-in and apical-out phenotype. (a) Immunofluorescent staining images of DAPI (blue), laminin-5 (red), and phalloidin (green) at days 16, 30, and 90 of their growth. Inset shows basal-in and apical-out phenotype with corresponding schematic. (b) Immunofluorescent staining images of DAPI (blue), Na^+^K^+^ATPase (red), and ezrin/villin-2 (green) at days 16, 30, and 90 of their growth. Inset shows basal-in and apical-out phenotype with corresponding schematic. All scalebars represent 100 µm, and scalebars in insets represent 50 µm.

Overall, the proposed mechanism for inversion in RPTEC organoids, wherein the addition of cold Matrigel to warm media with serum promotes partial gelling and orientation of the basal pole towards the organoid interior, was consistent with our previous reports of geometrically-inverted organoids [19, 20]. This interaction with Matrigel is the premise of the polarity reversal, and the basis for providing a soft scaffold to generate a 3D structure; rearrangement, cell movement, and rotation around the Matrigel ball could not be made possible in 2D culture. Such a mechanism highlights the importance of cell-material interactions for 3D culture to promote successful formation. It also underlines the importance of geometry, which has been of recent interest to the organoid community [33].

In our previous publication, we demonstrated applicability of the basal-in organoid platform with immortalized cells (MCF10A) [20]. In the current paper, we furthered this work to extended culture (90+ days) of primary proximal tubule cells. In comparison to immortalized MCF10A organoids, primary RPTEC organoids exhibited much more static growth patterns; in fact, RPTEC organoids decreased in size over time, compared to the increase in size of MCF10A organoids. Despite some morphological differences, both geometrically-inverted organoids showed good basal-in polarization, and maturation of the basement membrane over time, as indicated by laminin-5 immunostaining.

### 3.3. RPTEC organoid transcriptional profile

We further performed bulk RNAseq for the RPTEC organoids at day 31 of culture. All 3D cultures were compared to RPTECs grown in 2D well-plates, and all samples were run in triplicate. We generated volcano plots (Figure 4a) and found 3267 genes upregulated and 3278 genes downregulated. Furthermore, a heatmap (Figure 4b) was produced to compare gene expression for markers looking at transporters, EMT, maturity, and proliferation between 2D cultures and day 31 organoids. When comparing 2D cultures to day 31 organoids (Figure 4b), we observed upregulation of megalin (LRP2), aquaporin-1 (AQP1), Na^+^K^+^ATPase (ATP1A1), OCT2 (SLC22A2), MRP2, (ABCC2) MRP5 (ABCC5), MDR1 (ABCB1), MATE1(SLC47A1), and MATE2 (SLC47A2) in the organoids.

**Figure 4.**
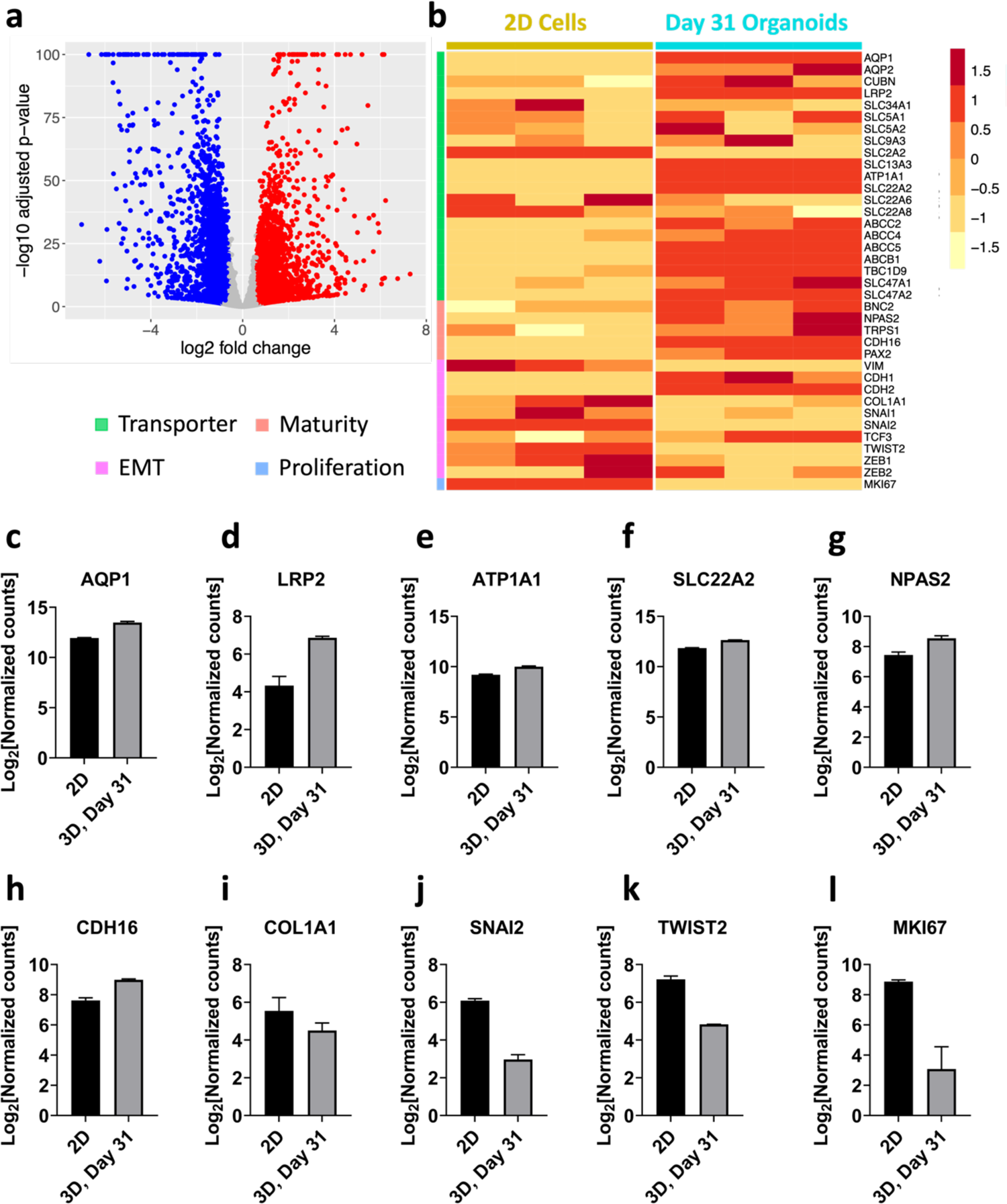
RPTEC organoid transcriptomics. (a) Volcano plot from bulk RNAseq comparing 2D RPTECs with day 31 organoids. (b) Heatmap from bulk RNAseq comparing 2D RPTECs to day 31 organoids. Normalized counts of genes in 2D RPTEC samples versus day 31 organoids for (c) AQP1, (d) LRP2, (e) ATP1A1, (f) SLC22A2, (g) NPAS2, (h) CDH16, (i) COL1A1, (j) SNAI2, (k) TWIST2, and (l) MKI67. N = 3 for both 2D and day 31 organoid samples. These genes were significantly different between 2D and 3D cultures based on the LRT with the Benjamini-Hochberg correction.

Next, we looked at genes related to EMT that are known to be increased in cells cultured on stiff substrates such as the typical 2D culture plastic. In comparison to 2D cultures on well-plates, the day 31 organoids had significantly lower expression for COL1A1, SNAI1, SNAI2, and TWIST2, all common markers for EMT. Hanging drop culture limits any interaction with the cells to surfaces of non-physiological elastic moduli, only media and the Matrigel scaffolding. Conversely, cultures maintained in 2D are grown on hard surfaces, which undergo EMT and preclude the possibility of long-term culture [10].These cell-matrix interactions and lack of EMT allow our basal-in and apical-out organoids to be sustained for long-term cultures (90 days), significantly different from 2D well-plate cultures. Furthermore, it is interesting to note that the organoids had higher e-cadherin and n-cadherin (Supplementary Figure 5); proximal tubule cells are one of the only epithelial cell types that express n-cadherin [34].

In terms of proliferation, it is a physiologically-relevant reflection of tubular maturation that MKI67 was downregulated in organoids compared to 2D cultures, an observation that was evident from the lack of growth and even shrinkage of the organoids during their culture period. Finally, we assessed more specific markers of a mature transcriptional profile [35, 36]. Compared to 2D well-plate culture, day 31 organoids had higher expression of NPAS2, CDH16, AQP1, LRP2, and PAX2.

We next plotted individual genes that were found to be significantly different genes, based on the likelihood ratio test (LRT) with Benjamini-Hochberg multiple comparisons corrections. A few genes were selected for transporters (Figures 4c-4f), maturity markers (Figures 4g-4h), EMT markers (Figures 4i-4k), and proliferation markers (Figure 4l). A more extensive list of significantly different genes can be found in Supplementary Figure 6. Taken together, these transcriptional results suggest that the organoids are in a mature and differentiated state, as they have downregulated proliferation compared to 2D cells, express increased maturity markers, and experience less EMT compared to 2D RPTECs.

### 3.4. Megalin (LRP2) and aquaporin-1 (AQP1) increase expression over time and in comparison to 2D culture

We aimed to explore indications of organoid maturity and differentiation through both immunofluorescence staining and bulk RNAseq. In line with the literature, we initially focused on aquaporin-1 (AQP1) and megalin (LRP2), as these particular markers are signs of cellular differentiation and maturation, something that is much more difficult to achieve in short-term 2D culture [10,17,18]. Immunostaining of AQP1 and LRP2 revealed an increased presence as the organoids matured. At day 16 of culture (Figure 5a), AQP1 and LRP2 staining are only present in certain cells. Not only does this indicate lack of full maturity, but also that the organoids exhibit some extent of cellular heterogeneity. As they were grown to day 90 (Figure 5b), a qualitative increase in both the percentage of AQP1 and LRP2 positive cells and signal strength can be observed.

**Figure 5.**
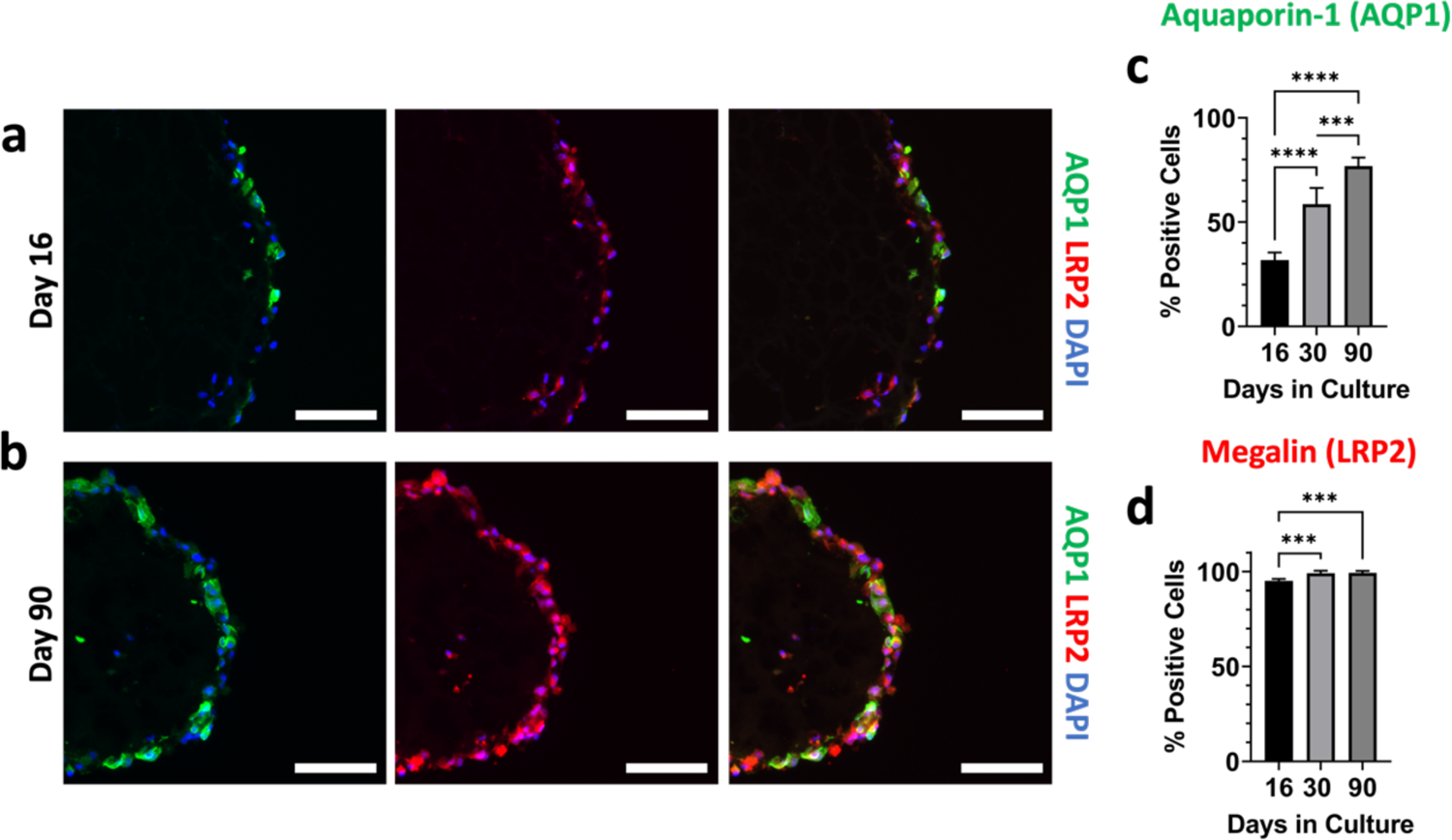
Organoids show maturity in comparison to 2D cultures and over 90 day growth period. Immunofluorescence images of aquaporin-1 (green), megalin (red), DAPI (blue), and the channels merged at (a) day 16, and (b) day 90 of their culture. All scalebars represent 100 µm. (c) Quantification of the percentage of cells that were positive for aquaporin-1 and (d) megalin. For panel (c) and (d), n=5 per timepoint. One-way ANOVA was performed using Tukey’s multiple comparisons. For both graphs, *** denotes p ≤ 0.001 and **** denotes p ≤ 0.0001.

In terms of quantitative measures, we counted the number of positive cells for both AQP1 and LRP2 at days 16, 30, and 90 of organoid culture. Immunofluorescent staining quantification revealed a significant increase in the percentage of AQP1 positive cells between all 3 days of culture (Figure 5c). Transcriptomics further revealed that AQP1 was higher for 3D culture compared to 2D well culture (Figure 4c). For LRP2, immunofluorescent quantification revealed that day 30 and 90 were significantly higher than day 16 (Figure 5d). It is important to note that nearly 100% of the cells at each timepoint were positive for LRP2. Normalized counts from sequencing showed that day 31 is significantly higher than counts from 2D cultures (Figure 4d). Taken together, these results indicate that RPTEC organoids: 1) increase expression of AQP1 and LRP2 over time and 2) express higher levels when compared to cells cultured on 2D well-plates. While both of these particular genes are of interest, we decided to focus on megalin because of its applications with protein uptake, proteinuria, and CKD.

In addition to the 3D nature of our cultures, another factor that may play a role in enhanced megalin expression is substrate stiffness, as is the case with inhibition of EMT. Previously, megalin has been shown to have higher expression on 3D polydimethylsiloxane (PDMS) constructs compared to 2D cells grown on stiff well-plates [37]. Similarly, another study showed that megalin expression is higher on 2D cells grown on 5 kPa PDMS compared to 50 kPa, underlining the importance of substrate stiffness [38]. While it is challenging to unravel the exact complexities, our results and previous findings in the literature suggest that substrate stiffness, cell-matrix interactions, and the 3D nature of our cultures all contribute to enhanced megalin expression.

### 3.5. Proximal tubule organoids exhibit time- and dose-dependent responses to proteinuric conditions

Because our proximal tubule organoid model: 1) has 3D architecture, 2) can be grown long term, 3) exhibits reversed polarization, 4) possesses consistently high megalin expression, and 5) increases megalin expression over time, we were motivated to use it for proteinuria studies. We were interested in such an application because proteinuria studies require apical access, since this is the surface on which megalin is expressed and protein reabsorption occurs, and long-term culture, as proteinuria is an important aspect of a chronic disease.

Before exposing kidney organoids to proteinuric conditions, we wanted to establish functionality of our organoids and confirm that the cells could uptake albumin. Our apical-out organoid model allows for this to be accomplished in an experimentally convenient manner, as the protein can be placed in the surrounding media, which represents the apical area. To do so, we incubated RPTEC organoids with fluorescein isothiocyanate bovine serum albumin (FITC-BSA) for 2 hours at 50 μg/mL, washed out excess FITC-BSA, and imaged the organoids using an in-incubator microscope (Incucyte S3, Sartorius). We observed FITC-BSA uptake for RPTEC organoids at day 16, 30, and 90 (Figure 6a and Supplementary Figure 6). Upon closely examining the GFP channel, smaller circular shapes can be observed corresponding to individual cells, suggesting successful uptake of the protein.

**Figure 6.**
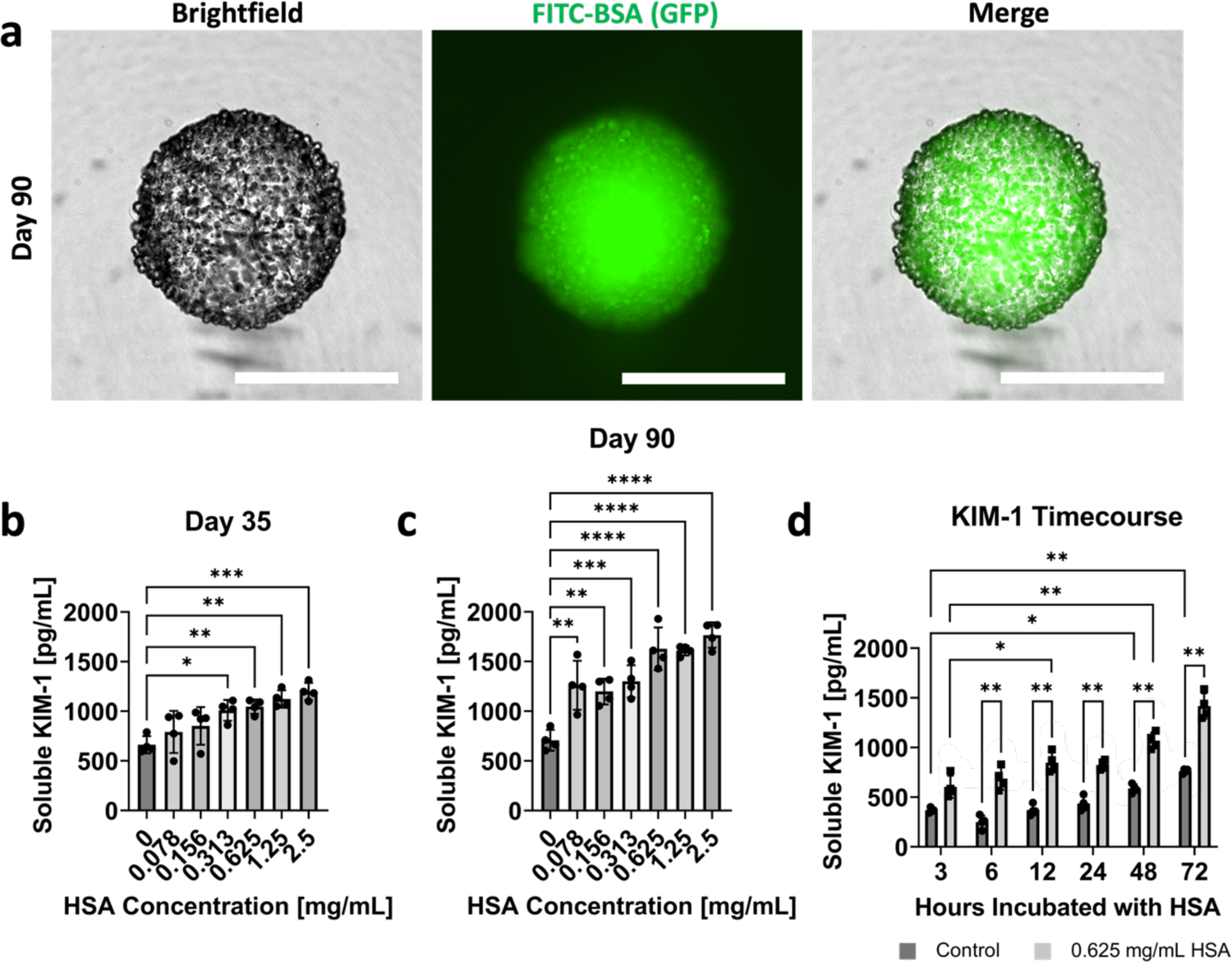
RPTEC organoids uptake albumin, and show dose- and time-dependent KIM-1 production to HSA stimulus. (a) Brightfield, GFP, and merged images of FITC-BSA uptake in day 90 RPTEC organoids. All scalebars represent 500 µm. Dose dependent KIM-1 production for organoids exposed to HSA at (b) day 35 and (c) day 90 of their culture. (d) Time-dependent KIM-1 production for control organoids and organoids exposed to 0.625 mg/mL HSA. For all plots, n = 4 was used for each dose or time point. One-way ANOVA with Tukey’s multiple comparisons was used for panels (b) and (c). Two-way ANOVA with Šīdák’s multiple comparison tests was utilized for panel (d). For all statistical tests, * indicates p ≤ 0.05, ** indicates p ≤ 0.01, *** indicates p ≤ 0.001, and **** indicates p ≤ 0.0001.

Upon confirmation of albumin uptake from the apical side of proximal tubule cells, we aimed to assess whether high protein levels can cause injury in the proximal tubule cells. We chose kidney injury molecule 1 (KIM-1) as a readout, as it is a non-invasive biomarker that is secreted into the culture media or urine. KIM-1 is a specific proximal tubule injury marker that has been widely established for nephrotoxicity and other kidney injury studies [5,9,39,40]. Conveniently, a portion of KIM-1 is secreted in the supernatant, making it easily detectable through an enzyme-linked immunoassay (ELISA). Consistent with the literature [41], we chose concentrations of HSA ranging from 0.078 mg/mL to 2.5 mg/mL in factors of 2, plus a control with no HSA present. First, we validated the ELISA kit with many acellular supernatant controls to ensure that no media components or growth factors caused false signal KIM-1 signal (data not shown). Organoids were then incubated with HSA in ULA plates for 48 hours, and the supernatant was collected at the timepoint. Our results show that the KIM-1 was produced in a dose-dependent manner for organoids exposed to HSA at day 35 (Figure 6b) and day 90 (Figure 6c) of culture. The KIM-1 data suggest that there is a more substantial dose dependent response with more mature organoids (day 90).

Since 0.625 mg/mL of HSA consistently showed a high response, we chose this condition to determine the time-dependence of KIM-1 production in day 35 organoids. Supernatant was collected from both control samples and 0.625 mg/mL HSA samples at 3, 6, 12, 24, 48, and 72 hour timepoints. The results indicated a time-dependent increase in KIM-1 cleavage for both the control and proteinuric conditions (Figure 6d). For a given condition, although each timepoint is not significantly different from one another, there is an overall significance of the increase in KIM-1 production. Furthermore, at each of the timepoints, the KIM-1 production was significantly higher for the proteinuric condition compared to the control. Because of the dose- and time-dependent responses from HSA exposure seen in KIM-1 production, this geometrically-inverted proximal tubule organoid shows promise for future proteinurea studies. It also opens the possibility of studying nephrotoxictiy of drugs such as gentamycin and cisplatin, both of which have been implicated in apical uptake via megalin [42, 43].

## 4. Conclusions

This study establishes the first geometrically-inverted (basal-in, apical-out) proximal tubule model, to our knowledge, using normal RPTECs. From a biomaterials perspective, the results suggest a broader utility of our high-throughput minimal Matrigel scaffolding technique, originally used with a mammary epithelial cell line, to primary human epithelial cells such as from the renal proximal tubule. The maintenance of the proximal tubule organoids for 90+ days is important biologically for enhanced tissue maturation. From a technological perspective this extends, by two months, the demonstrated limit of duration of continuous hanging drop culture. More specifically, immunostaining revealed biomaterial-induced polarity inversion of the normal epithelial cell-derived organoids, as apical markers such as phalloidin and ezrin/villin2 were located on the organoid exterior (apical-out), and basolateral markers such as laminin-5 and Na^+^K^+^ATPase were located on the organoid interior (basal-in). RNAseq revealed that day 31 organoids show less EMT and proliferation compared to 2D-cultured RPTECs, while they have increased levels of maturity markers and transporters. Focusing on proteinuria as a future application enabled by geometrically-inverted organoids, we analyzed functional protein uptake and response. Indeed, fluorescent-albumin was readily taken up from the media surrounding the organoids, and albumin-induced injury, as measured by the early injury marker KIM-1, was observed in a dose- and time-dependent manner. The important endocytic receptor for proteins, megalin, had higher mRNA expression in our organoids compared to 2D-cultured cells, and exhibited increased protein expression over time. Although this paper focuses its functional analysis on albumin-uptake and injury, the ease of apical access together with the generally high levels of transporter and endocytic receptor expression suggests broader utility, including certain types of drug-induced nephrotoxicity, where apical exposure is involved.

## Credit author statement

EP, JHL, and AYL: Conceptualization, investigation, methodology, writing, formal analysis, and visualization. XZ and ST: Conceptualization, methodology, investigation, writing, and supervision.

## Data availability

All data is available from the corresponding author upon reasonable request. Our RNAseq data have been deposited on Gene Expression Omnibus (GEO) of NCBI under accession number GSE198857.

## Acknowledgements

This work was supported by the National Institutes of Health grant number SC1DK112151 and the Price Gilbert Jr. Chair Fund. This study was supported in part by the Emory Integrated Genomics Core (EIGC), which is subsidized by the Emory University School of Medicine and is one of the Emory Integrated Core Facilities. We thank Professor Greg Gibson and the Georgia Tech Center for Integrative Genomics for advice on bioinformatics analysis. This research was supported in part through research cyberinfrastructure resources and services provided by the Partnership for an Advanced Computing Environment (PACE) at the Georgia Institute of Technology, Atlanta, Georgia, USA. This material is also based upon work supported by the National Science Foundation Graduate Research Fellowship Program to EP (Grant Number: DGE-1650044). All schematics in this paper were generated using Biorender.

## Supplementary Materials Methods

### Live/dead stain

The Live/Dead Viability/Cytotoxicity Kit (Invitrogen, #L3224) was used to assess organoid viability. Samples were harvested from hanging drop plates, washed with PBS 3 times, and incubated with the live/dead solution for 20 minutes at room temperature. The staining solution consisted of 2 µM Calcein-AM (live stain) and 4 µM ethidium homodimer-1 (dead stain) in PBS. Samples were immediately imaged after the incubation period using a DMi8 epifluorescence microscope (Leica). In the images, green denotes live cells and red denotes dead cells.

### Statistical analysis

Area, diameter, and roundness were quantified for two independent experiments. For the first one (Supplementary Figures 2a-2d), 31 organoids were imaged on day 90. For the second independent experiment, 22 organoids were tracked from days 77 to 103 (Supplementary Figures 2e-2i). No statistical analysis was performed.

## Supplementary Figures

**Supplementary Figure 1.**
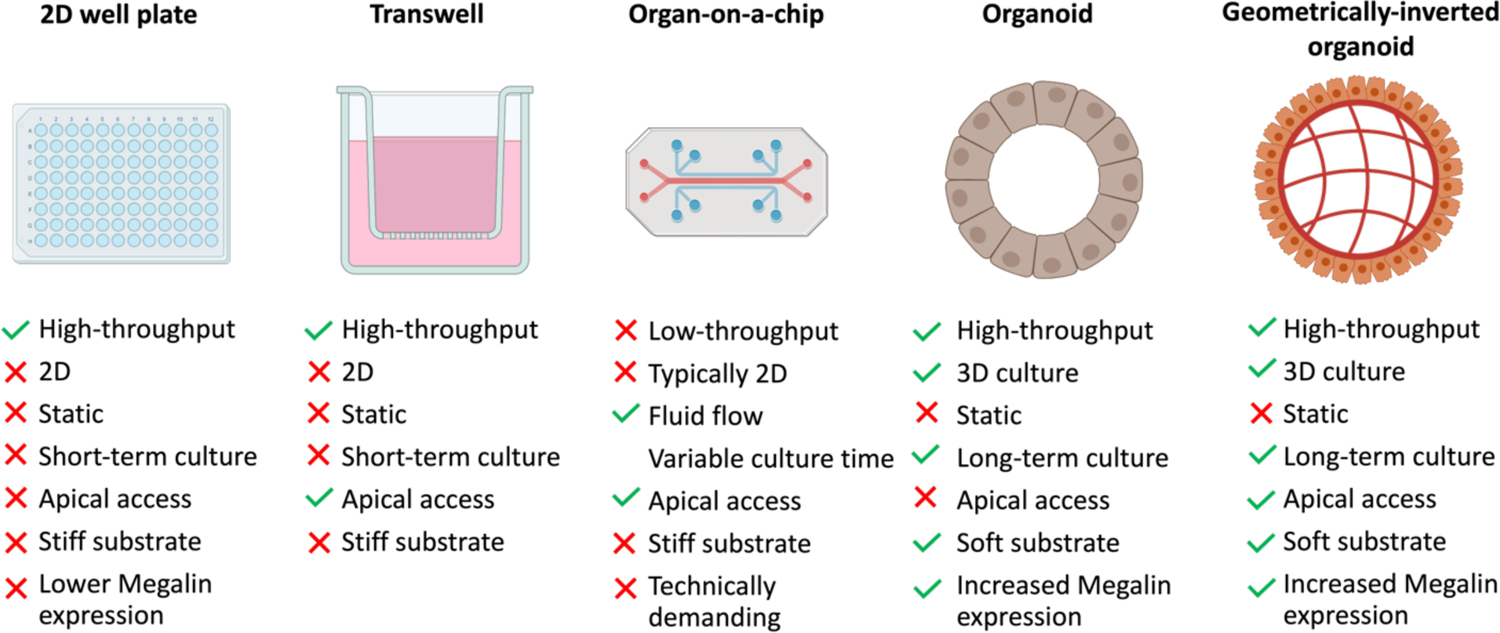
Summary of existing *in vitro* models for proteinuria and chronic kidney disease (CKD) studies and our proposed geometrically-inverted organoid, with advantages and disadvantages of each highlighted.

**Supplementary Figure 2.**
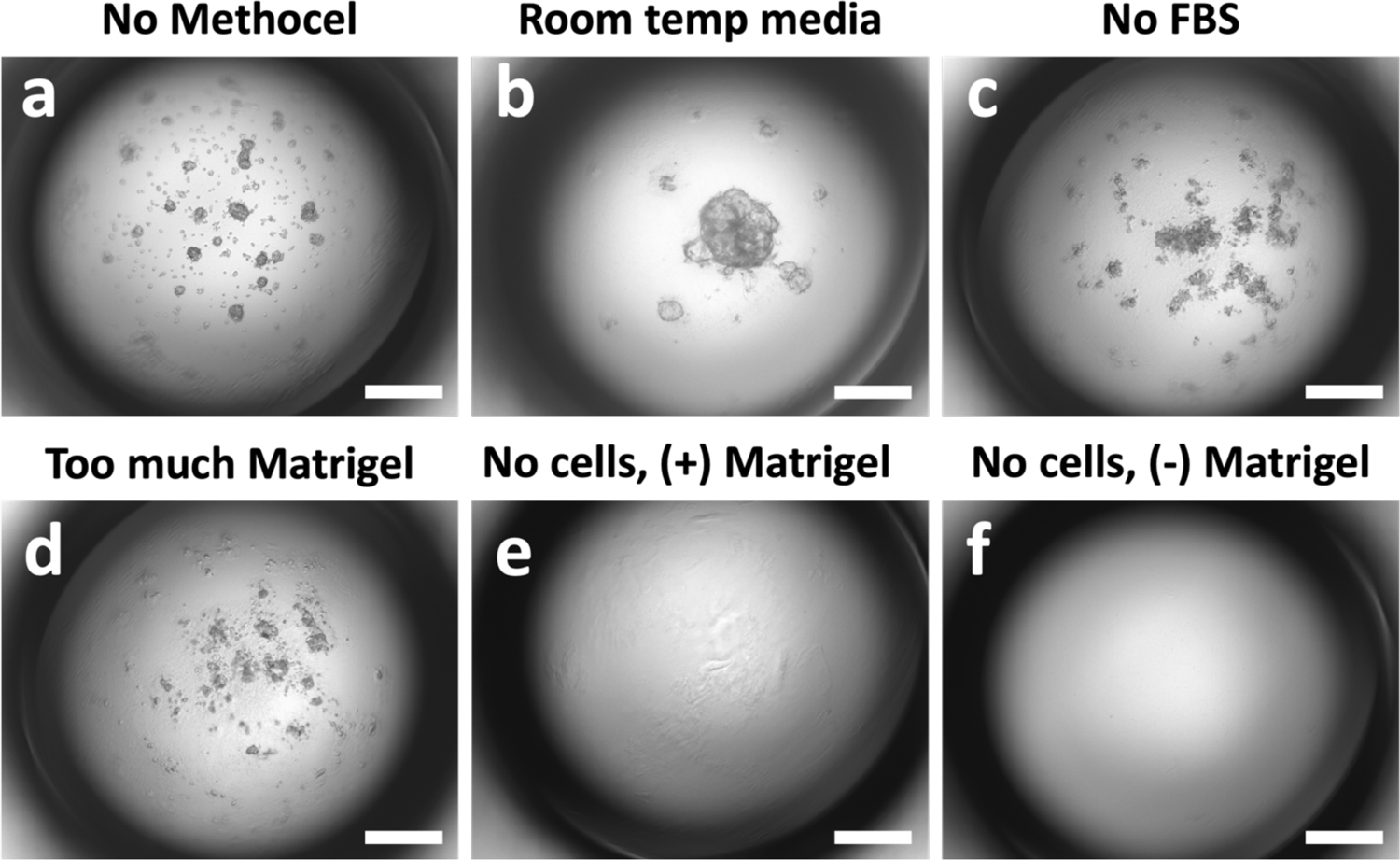
Optimization of organoid seeding and minimal Matrigel Scaffold. Improper organoid formation when (a) Methocel was not included, (b) Matrigel was added to room temperature mixture of growth media, FBS, and methylcellulose, (c) no serum was included in the seeding media, and (d) too much Matrigel was included in the seeding media. Acellular seeding media consisting of growth media, FBS, and Methocel (e) with the addition of Matrigel, and (f) without the addition of Matrigel. All scalebars represent 500 µm.

**Supplementary Figure 3.**
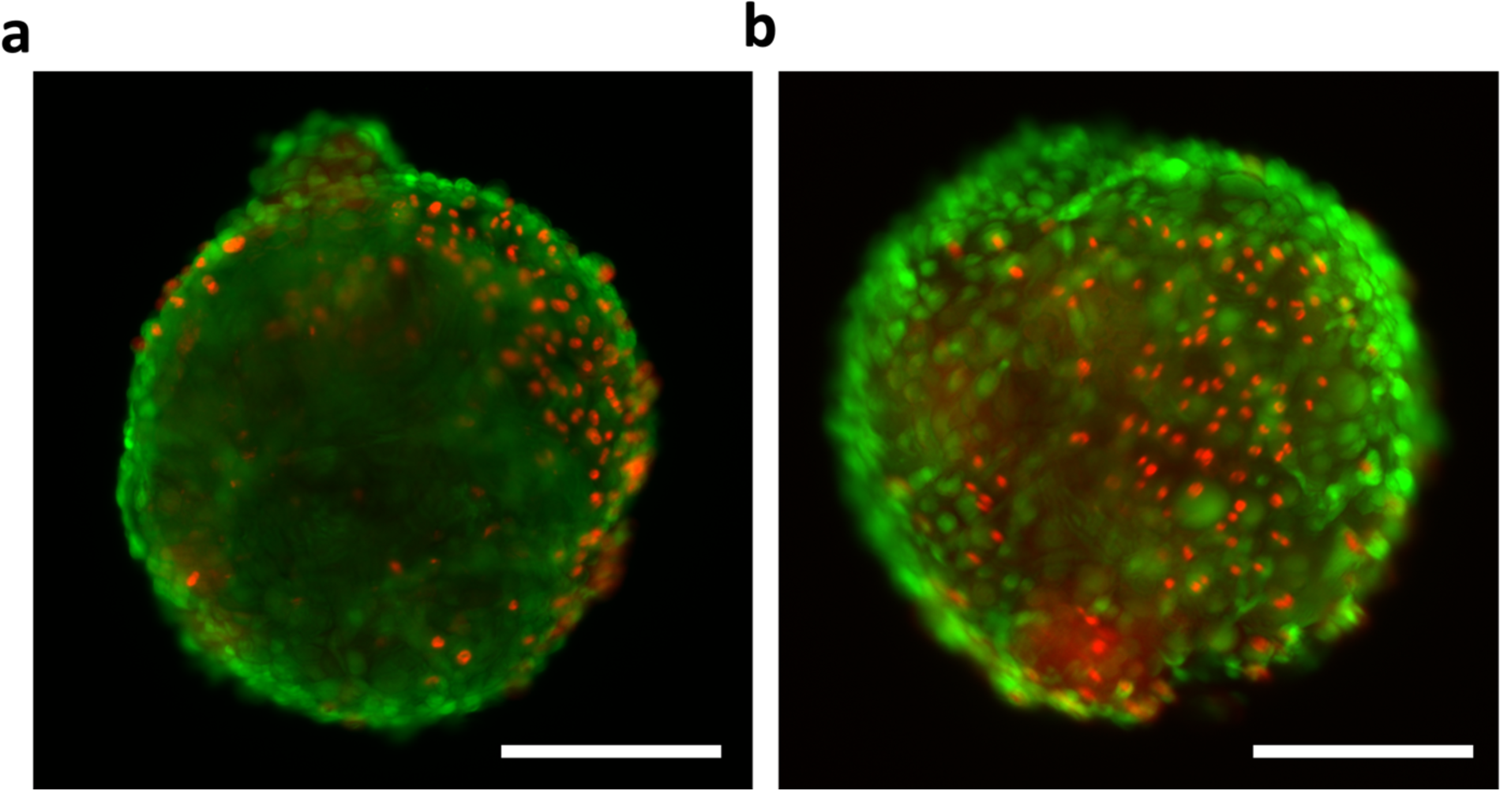
RPTEC organoids remain viable during 90-day culture period. Live/dead staining of RPTEC organoids at (b) day 65 and (c) day 90 of their culture. Green represents live cells, and red represents dead cells. Scalebars represent 250 µm.

**Supplementary Figure 4.**
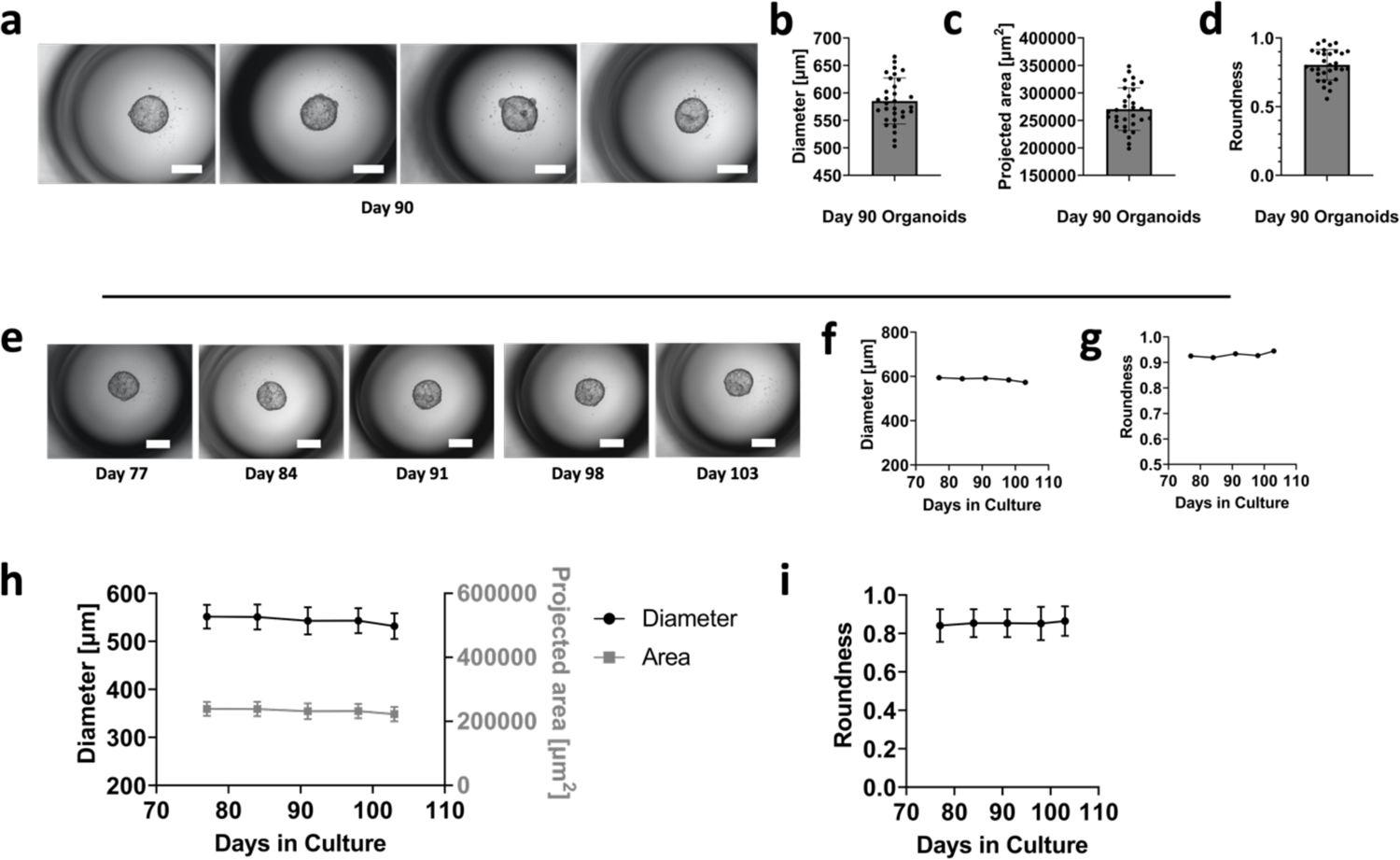
Extended morphology analysis from 2 independent experiments. (a) Representative images of day 90 organoids from an independent experiment. (b) Diameter, (c) projected area, and (d) roundness of day 90 organoids, n = 31. All scalebars represent 500 µm. (e) Image of organoids tracked over time from an independent experiment. (f) Diameter, and (g) roundness values for the organoid shown in (e). (h) Average diameter and (i) average roundness of n = 22 organoids per timepoint.

**Supplementary Figure 5.**
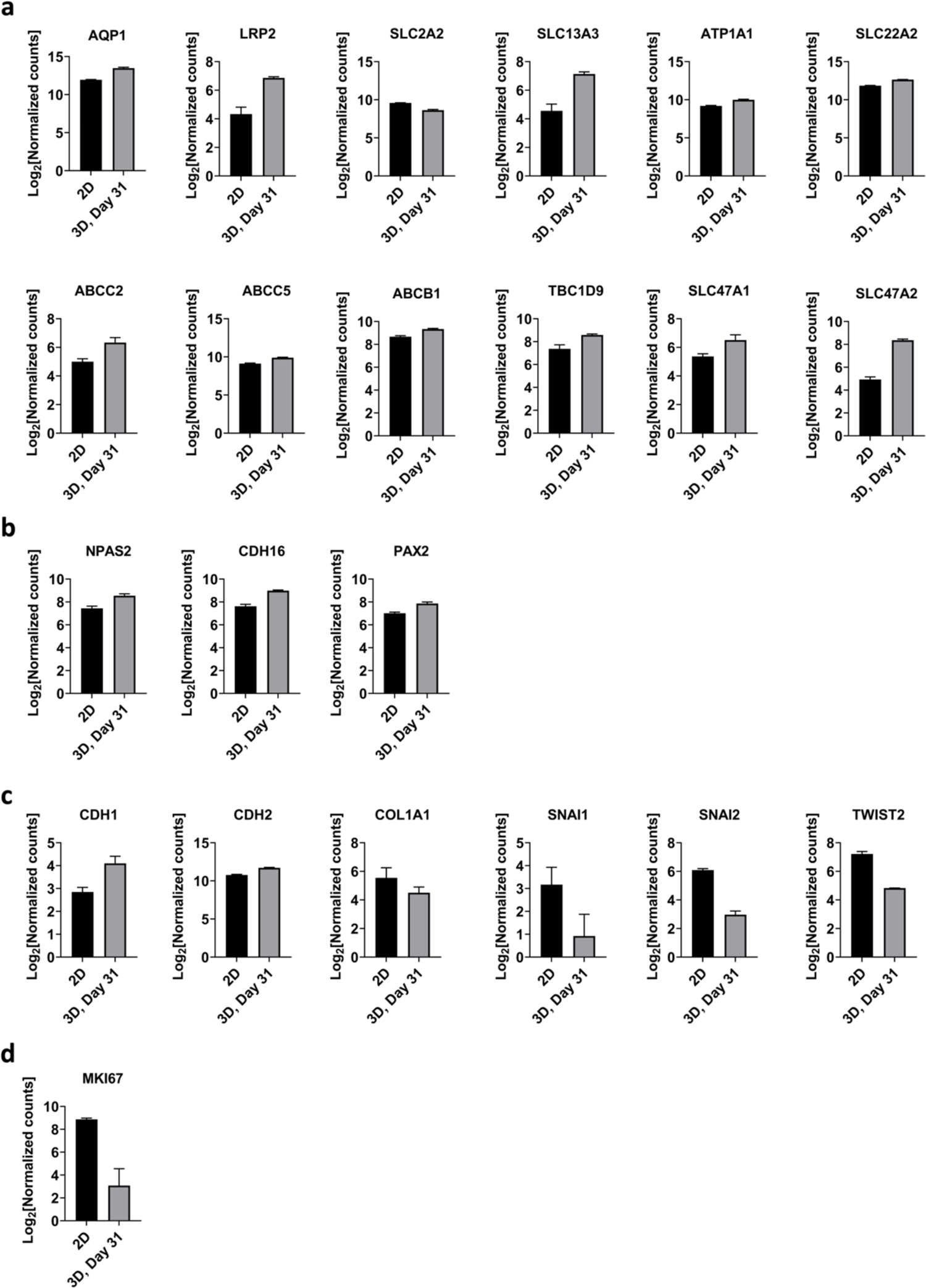
Transcriptional analysis of significantly different genes. Comparison of gene expression levels for markers related to (a) transporters, (b) maturity, (c) EMT, and (d) proliferation. All samples were run in triplicate. All genes were significantly different between 2D and 3D cultures based on the LRT with the Benjamini-Hochberg correction.

**Supplementary Figure 6.**
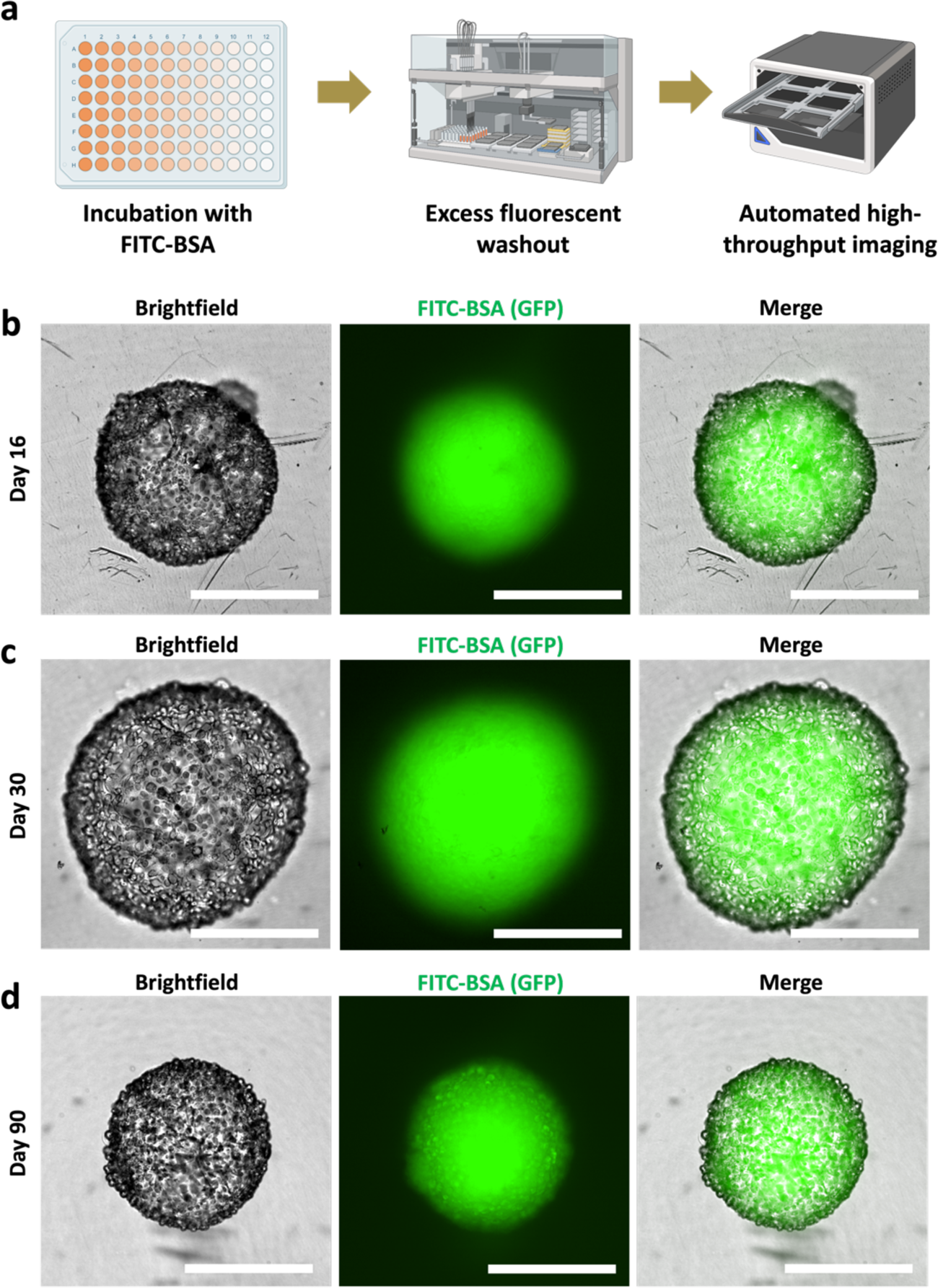
RPTEC organoids successfully uptake FITC-BSA. (a) Schematic of procedure used for albumin uptake assay. Brightfield, GFP, and merged images of FITC-BSA uptake in (b) day 16, (c) day 30, and (d) day 90 RPTEC organoids. All scalebars represent 500 µm.

